# MetaLLM: Residue-wise Metal ion Prediction Using Deep Transformer Model

**DOI:** 10.1101/2023.03.20.533488

**Authors:** Fairuz Shadmani Shishir, Bishnu Sarker, Farzana Rahman, Sumaiya Shomaji

## Abstract

Proteins bind to metals such as copper, zinc, magnesium, etc., serving various purposes such as importing, exporting, or transporting metal in other parts of the cell as ligands and maintaining stable protein structure to function properly. A metal binding site indicates the single amino acid position where a protein binds a metal ion. Manually identifying metal binding sites is expensive, laborious, and time-consuming. A tiny fraction of the millions of proteins in UniProtKB – the most comprehensive protein database – are annotated with metal binding sites, leaving many millions of proteins waiting for metal binding site annotation. Developing a computational pipeline is thus essential to keep pace with the growing number of proteins. A significant shortcoming of the existing computational methods is the consideration of the long-term dependency of the residues. Other weaknesses include low accuracy, absence of positional information, hand-engineered features, and a pre-determined set of residues and metal ions. In this paper, we propose MetaLLM, a metal binding site prediction technique, by leveraging the recent progress in self-supervised attention-based (e.g. Transformer) large language models (LLMs) and a considerable amount of protein sequences publicly available. LLMs are capable of modelling long residual dependency in a sequence. The proposed MetaLLM uses a transformer pre-trained on an extensive database of protein sequences and later fine-tuned on metal-binding proteins for multi-label metal ions prediction. A 10-fold cross-validation shows more than 90% precision for the most prevalent metal ions.

## Background

Proteins are biomolecules composed of amino acid chains that form the building blocks of life, and play fundamental roles in the entire cell cycle. They perform multitudes of functions including catalyzing reactions as enzymes, participating in the body’s defense mechanism as antibodies, forming structures, and transporting important chemicals. In addition, they interact with other molecules including proteins, DNAs, RNAs, and drug molecules to act on metabolic and signaling pathways, cellular processes, and organismal systems. Protein structures and interactions describe the molecular mechanism of diseases and can convey important insights about disease prevention, diagnosis, and treatments. Likewise, proteins bind to different metal ions, such as zinc, iron, copper etc. to play necessary roles in many biological processes, including enzyme catalysis, regulation of gene expression, and oxygen transport. Metal ions are often bound to specific sites on proteins, known as metal binding sites, which play a key role in determining protein’s structure and function. Identifying metal binding sites manually using various experimental procedures such as mass spectrometry, electrophoretic mobility shift assay, metal ion affinity column chromatography, gel electrophoresis, nuclear magnetic resonance spectroscopy, absorbance spectroscopy, X-ray crystallography, and electron microscopy is an expensive, laborious, and time-consuming process [21]. A very small fraction of the millions of proteins stored in UniProtKB [1] – the most comprehensive protein database – are annotated with metal binding sites. Millions of other proteins are awaiting for metal binding site annotation. To keep pace with the exponential increase of protein sequences in the public databases, it is essential to develop computational approaches for predicting metal binding sites in proteins. Considering the benefits it can provide in understanding function and structure of proteins as well as having practical implications in drug design and biotechnology, automatic prediction of metal binding sites in proteins is considered to be an important problem in computational biology.

Predicting the binding sites for metals is a challenging problem in computational biology. Decades of research has been dedicated to discovering computational approaches that can accurately predict the metal ions as well as the positions where they bind to the proteins [3, 17, 8, 5]. A comprehensive review of recent advances in computational approaches for predicting metal binding sites can be found in [30]. Broadly, these approaches can be categorized into following three groups based on the type of attributes they take into account: 1) structure-based methods that use three dimensional secondary structure of proteins as primary data; 2) sequence-based methods that use amino acid sequence as primary data; and 3) combined methods that leverage both structure and sequence attributes.

Structure-based approaches for predicting metal binding sites use a combination of geometric, chemical, and electrostatic criteria to explain the metal-protein interaction that eventually works to identify possible binding sites in a protein structure. A relatively early work described in [26] uses electrostatic energy computation [20, 9] to find binding affinity of metal ion to a site in a protein structure. In [4], the proposed method learns geometric constraints to differentiate binding sites for different metals based on the statistical analysis of structures of metal binding proteins. Another structured-based method is proposed in [33] for predicting only the zinc binding sites. A template-based method is proposed in [19] where a database of pre-computed structural templates for metal binding sites is searched against each residue in a query protein to find which metal binds to it. mFASD [12] is a structure based model to predict metal binding sites. From the structures of metal binding proteins, mFASD computes the functional atom set (FAS) - the set of the atoms that are in contact with the metal - for each metal, and store it as reference. The distance between FASs of different metals is used to distinguish binding sites for different metals. Given a query protein structure, mFASD scans the database reference FASs against FAS of each sites, and computes the distance. The decision is made based on how many reference FASs matched for a given metal. One of the shortcomings of the structure based models is that they are dependent on structural databases such as Protein Data Bank (PDB) which is very limited in terms of amount of protein structures in the database.

On the other hand, sequence-based methods for predicting metal binding sites use sequence conservation, alignment, and similarity to identify metal binding sites. For example, [2] is a sequence-based method that find the patterns of binding sites from the metal binding proteins. Sliding window-based feature extraction and biological feature encoding techniques are proposed to predict the protein metal-binding amino acid residues from its sequence information using neural network in [15] and using support vector in machine [16]. MetalDetector [18] is a sequence-based technique that uses decision tree to classify histidine residues in proteins into one of two states as 1) free, or 2) metal bound; and cysteine residues into one of three states as 1) free, 2) metal bound, or 3) disulfide bridged. A two stage machine learning model is proposed in [22] that includes support vector machine as local classifier in the first stage, and a recurrent neural network (RNN) [13] in second stage to refine the classification based on dependencies among residues. A combined approach proposed in [27] where support vector machine, sequence homology, and position specific scoring matrix (PSSM) are put together to predict zinc-binding Cys, His, Asp and Glu residues.

Additionally, there are combined methods for predicting metal binding sites that use an ensemble of sequence, structural, and physicochemical features. For example, MetSite [28] is a method that uses both the sequence profile information and approximate structural data - PSSM scores together with secondary structure, site residue distances, and solvent accessibility - are fed into neural network machine learning technique.

Machine learning techniques such as decision tree, random forests, support vector machine, neural network have been widely applied to the problem of metal binding site prediction. Recently, deep learning based techniques such as convolutional neural network (CNN) [14], long short term memory (LSTM) [13], transformers [29] etc. are increasingly used in metal binding site prediction to leverage huge amount of sequence data deposited into the protein databases. These advanced models are very efficient in handling large datasets. For example, [10, 11] applied different deep learning architectures namely 2D CNN, LSTM, RNN coupled with various feature extraction techniques for predicting metal-binding sites of Histidines (HIS) and Cysteines (CYS) amino acids. Classical machine learning techniques working on protein sequences mostly depends on hand-engineered features and can not model long-distance residual dependency. Moreover, fitting these models on large datasets is a big challenge.

Many of the existing methods are performing with good accuracy. However, predicting metal binding sites is a challenging problem because it is difficult to identify and distinguish the presence of different metals with similar chemical properties. A major shortcoming of the existing computational methods is in taking into account the long-distance dependency of the residues to distinguish the presence of distinct metal ions. There are other shortcomings such as low accuracy, absence of positional information, hand-engineered features, and pre-determined set of residues and metal ions. Building high performing prediction model would require a comprehensive understanding of the structural, chemical, and biological factors that influence metal binding. Therefore, ongoing research effort is important to improve our understanding of metal binding in proteins and to develop more accurate and efficient computational pipeline to keep pace with the growing number of proteins.

Considering the challenges and the availability of protein data, in this paper, we propose MetaLLM, a metal and binding site prediction technique by leveraging the recent progress in self-supervised attention-based (e.g. Transformer) large language models (LLMs) and huge amount of protein sequences publicly available. LLMs are capable of modeling long residual dependency in a sequence. The proposed MetaLLM uses a transformer pre-trained on large database of protein sequences, and later fine-tuned on metal binding proteins for multi-label metal ions prediction In the fine-tuning step, the low rank sequence embeddings are concatenated with positional one-hot embeddings and fed into the fully connected neural network layer for multi-label metal ions prediction. A 10-fold cross-validation shows more than 90% precision for most prevalent metal ions.

### Methodology

In this section the proposed methodology and overall workflow has been discussed thoroughly. The process mainly consists of four major steps: (i) using a transformer-based model to generate prominent numerical embeddings from raw protein sequence, (ii) generating the position embedding of the metal ions to locate the site for binding, (iii) concatenating sequence and position embedding together to find a combined vector which consists of prominent features from the sequence along with the site for binding, and (iv) considering the combined vectors as training sample, trained a neural network for predicting metal ion candidates which will be bound to a particular site. An overview of the individual concepts are presented first, and then the proposed methodology is discussed in details. The notations of the proposed model are shown in **Table** 1.

**Table 1.** Notations and their descriptions.

| Notation | Description |
| --- | --- |
| $x$ | input |
| $y$ | output |
| $n$ | number of samples |
| $\eta$ | learning rate |
| $\sigma$ | activation function |
| $\theta$ | set of parameters |
| $d_k$ | queries of transformer |
| $d_v$ | key-values of transformer |
| $BCE$ | Binary cross entropy |

### Definitions and Notations

**D**eep Learning is a machine learning technique that trains computers to perform what might be natural to humans: learning by example. Deep learning models are built using neural networks, which are inspired by the structure and function of the brain [14]. Layers of interconnected nodes process vast amounts of complex data, making deep learning excel at learning from the experience. Due to the prominent feature extraction ability, deep learning is considered as one of the most powerful and robust classifier tools and is a key technology behind numerous challenging applications. For instance, deep learning is integrated into autonomous vehicles-enabling them to detect a road sign, or distinguish a pedestrian from a car. It has also a great impact on business, government, and tech companies.

As mentioned above, deep learning, also known as representation learning, is a promising feature extraction method that allows an algorithm to be fed with raw signals *x* (e.g., pixels of an image), generates their internal representations in such a way that is useful for regression or classification for similar data, and finally utilizes the learnt representation to predict the the class label *y* of an unseen data. By leveraging the non-linear activation function, the diversity among the training data are learnt and distinguished better to achieve the state-of-the-art classification outcomes.

**T**ransformer is a neural network architecture primarily used for natural language processing tasks, e.g., language translation or text summarization etc. The transformer architecture was introduced in the paper [29]. One of the main advantages of the transformer architecture lies on its ability to process input sequences in parallel, rather than in a sequential manner like many other models. This makes it well-suited for tasks that involve long input sequences and consequently speed up the training and inference process. Similar to other neural networks, the transformer architecture has also achieved state-of-the-art results on many natural language processing tasks. Generally, a transformer has an encoder and a decoder neural network. It basically follows the structure using layers of self-attention, fully connected layers for both the encoder and decoder architecture, respectively. The input of the encoder consists of queries as well as keys of dimension *d*_*k*_, and values of dimension *d*_*v*_. There are dot products of the query with all keys, which will divide each by 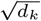, and apply a softmax function to obtain the parameters. The attention function on a set of queries was done simultaneously, packed together into a matrix Q. The keys and values are also packed together into matrices K and V. The outputs as:

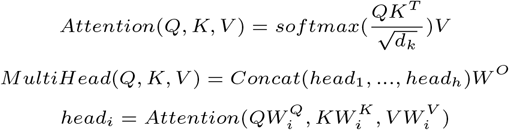

### MetaLLM pipeline

In this section, some relevant concepts related to our proposed model architecture are discussed. Firstly, the transformer-based language models are introduced and then the discussion on how the protein site prediction problem can be deciphered with the language models together, is presented. In recent years deep learning and machine learning have been advancing with many domains like computer vision and natural language processing (nlp). These breakthroughs happened with the help of powerful algorithms and high performing hardwares. Algorithms, like transformers and auto-regressive models, leverage not only nlp, but also many aspect of computational biology as well. Recent studies show that transformer based model are using heavily in both academia and industry for various life science experiments, like protein folding, drug discovery, and gene expression prediction [31]. Like other deep learning models, ProtTrans is a transformer based model, including attention mechanism and positional encoding, was trained on tons of data from different data sources containing up to billions of amino acid. In order to capture the biophysical features of the proteins, the dimensionality reduction technique was used to visualize the prominent features for subsequent supervised task. By leveraging transfer learning for the downstream task, ProtTrans was considered as a potential candidate for our methodology, since, transfer learning benefits to do prediction and detection with limited amount of data using the pre-trained weights of the model. By using this methodology, one can easily train deep learning model for a very specific task with a very low computational cost.

### Proposed Model Architecture

Our proposed model in **Figure 1** initially employs a pre-trained language model called ProtTrans which effectively generate embedded and informative protein sequence representation from the raw protein sequence. ProtTrans, a state-of-the-art protein language model, was used as the back-end of our proposed architecture. Moreover, we have also tested our representation for the classification of various metals to justify the novelty of the language model. Our proposed architecture leverages the state-of-the-art ProtTrans model for the feature extraction, which was trained on 5616 GPUs and TPUs Pod up-to 1024 cores [7]. Then, the embedded 1024-dimensional features were extracted from the last layer of ProtTrans model and classification was done integrating the sites information with the protein sequence embedding. The fixed 1024 dimension was chosen through the investigating and most discriminant features Our novel contribution on this paper is to carefully design a neural network which was more robust and cost effective with limited number of parameters without dropping the accuracy of the prediction. The metal prediction was done by the classification model on top of the language model. The input of the classification layer was the 1024 size embedding from the language model and then concatenated with the same dimensional sites information. The weights of the four layers of classification model was initialized randomly and for the non-linearity, relu activation function was used. In order to get the final prediction, sigmoid activation function was used since it is a multi-class multi-label prediction. We also used 10-fold cross validation to evaluate the whole dataset better and maximize the performance of our model. The mathematical representation are given in equations respectively.

**Fig. 1.**
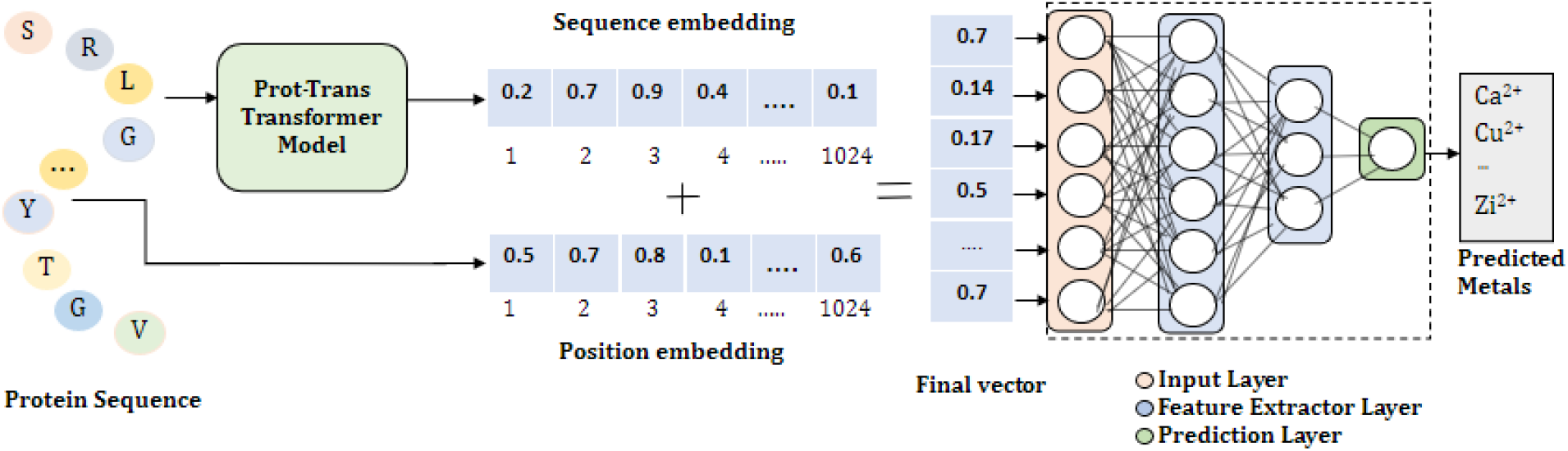
The overall architecture of the proposed framework of MetaLLM: (i) the protein sequences are delivered as an input to the pre-trained language model to produce the sequence embedding, (ii) the embedding for identifying the binding positions of the amino acids are extracted (ii) both the embedding are concatenated to obtain a final vector, and (iv) the final vectors are fed into a neural network to predict the metal ions.

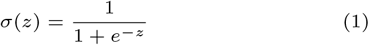

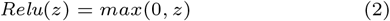

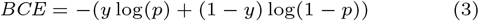

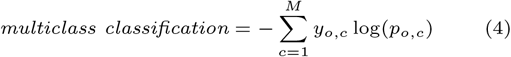

## Results and Discussion

### Dataset Description

In this work, we prepared a dataset of metal binding proteins extracted from UniProtKB [1]. We have extracted the proteins that are annotated with presence and binding site of metal ions. The protein sequences are further filtered to keep only those sequences with a length in the range of 50 to 1000 amino acids. However, some metals have limited number of samples. Those samples were removed during the pre-processing step. The sample distribution of the six metals are shown in **Figure 2** After necessary pre-processing, the final dataset contains 18,348 number of protein sequences holding 6 number metals in total.

**Fig. 2.**
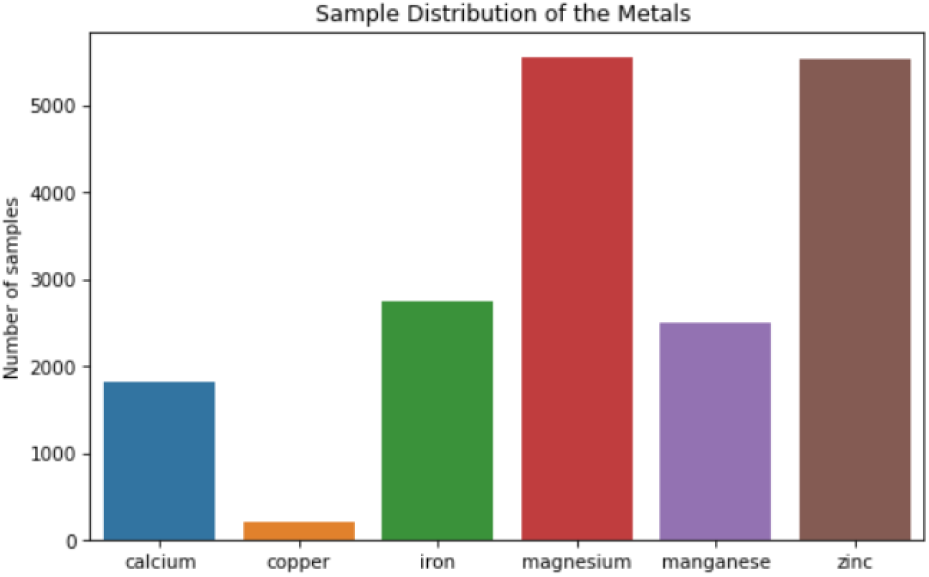
This figure shows the number of samples of our dataset.

### Implementation details

MetaLLM is implemented using Bio-Transformers^1^ python package that wraps the implementation of ProtTrans (Protein Transformer) [7] and ESM (Evolutionary Scale Modeling) [25, 24, 23] - two of the large language models trained on million of protein sequences to predict embeddings. MetaLLM uses the ProtTrans as the backend model which is pre-trained on 393 billion amino acids. Pre-trained models are already trained on a large dataset using expensive devices such as numerous powerful GPUs to learn the vector representation of sequences. These models then can be readily used to get low rank vector representation also known as embeddings given a new sequence. As a first step, MetaLLM gets the sequence embeddings usign ProtTrans backend. Alongside, the positions of the binding residues are encoded using multi-hot embeddings. These two embeddings 1) sequence embeddings, and 2) Position embeddings are combined together to get a resultant vector representing metal binding protein sequence and the binding sites. This resultant vector is then feed into the fully connected neural network to train a model end-to-end for predicting metal ions that bind the encoded positions. The proposed model is consist of four-layer fully connected neural network with 500 hidden neurons in the first layer, 300 neurons in the second and third layer, and final layer with 100 hidden neurons before it goes to soft-max layer for predicting the probability of different metals. To prevent the over-fitting of the network, we carefully used a dropout with a rate of 0.02. Different batch sizes were investigated during the experiment, and 64 is found to be the optimum batch size. While selecting the learning rate in our training, we used a range from 0.0001 to 0.01 for evaluating the results. For the optimization, the state-of-the-art adam optimizer was used with the binary cross entropy loss function. Early stopping was also used to make the model stable during training. The experiment was done on Nvidia GeForce RTX 3090 GPU, and the training phase took 45 minutes to complete the 200 epochs. During the test phase, it took approximately 5 seconds to make the predictions for one batch of protein sequence. The chosen hyper-parameters of the best proposed architecture are listed in **Table** 2. During the evaluation phase, we have run a 10-fold cross-validation, and the results are reported in terms of precision, recall and f1-score.

**Table 2.** Hyper-parameters of the Model

| Hyper parameter | value |
| --- | --- |
| Learning rate | 0.001 |
| Batch size | 64 |
| Epoch | 200 |
| Dropout | 0.02 |
| Momentum | 0.9 |
| Number of layers | 4 |
| Neurons in last layer | 100 |

### Evaluation Metrics

To evaluate the performance of our proposed system, four long-established evaluation metrics for measurement have been considered, they are: Recall, Precision, f-measure (f1), and Accuracy. The equations for the metrics are following:

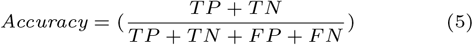

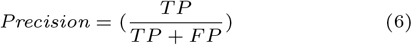

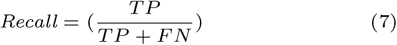

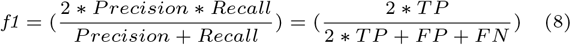

In the above mentioned equations, true positives (TP) and true negatives (TN) denote the number of correctly predicted metals, and false positives (FP) and false negatives (FN) denote the number of incorrectly predicted metals.

## Results

We evaluated MetaLLM by Precision, Recall, and f1-score using the 10-fold cross-validation (CV) and independent test sets of different metals with and without positional information. As shown in **Table 3**, the MetaLLM model obtained highest precision value for magnesium of 0.97 and highest recall value of 0.94 for iron and magnesium on the 10-fold CV. On the other hand, the highest f1-score was obtained of 0.95 for iron and magnesium respectively. These results indicated that stabilizing the ProtTrans language model may potentially capture the evolutionary and structural information of the protein with the help of positional information. We also conducted our experiments without having the positional information. In **Table 4**, it is shown that the precision, recall, and f1-scores were slightly dropped when the positional information was not concatenated with the protein embedding. MetaLLM achieved validation accuracy of 90% when the position of the metals was concatenated. The bar plot of the different metrics are shown in **Figure 3, Figure 4**, and **Figure 5**. To compare our accuracy of the model with other models, we also did an experiment developing a multi-label k-nearest neighbours [32]. And, we got 73% of validation accuracy while our proposed model had 90% of validation accuracy. **Figure 6** shows the training and validation accuracy of our proposed model.

**Table 3.** Performance metrics of the proposed model (with positional information)

| Metal Name | Precision | Recall | f1-score | Support |
| --- | --- | --- | --- | --- |
| calcium | 0.83 | 0.84 | 0.84 | 202 |
| copper | 0.94 | 0.67 | 0.78 | 24 |
| iron | 0.96 | 0.94 | 0.95 | 331 |
| magnesium | 0.97 | 0.94 | 0.95 | 47 |
| manganese | 0.92 | 0.93 | 0.92 | 243 |
| zinc | 0.91 | 0.89 | 0.90 | 558 |

**Table 4.** Performance metrics of the proposed model (without positional information)

| Metal name | Precision | Recall | f1-score | Support |
| --- | --- | --- | --- | --- |
| calcium | 0.94 | 0.71 | 0.81 | 279 |
| copper | 0.67 | 0.93 | 0.78 | 15 |
| iron | 0.89 | 0.90 | 0.90 | 100 |
| magnesium | 0.72 | 0.94 | 0.82 | 433 |
| manganese | 0.71 | 0.47 | 0.56 | 326 |
| zinc | 0.84 | 0.95 | 0.89 | 616 |

**Fig. 3.**
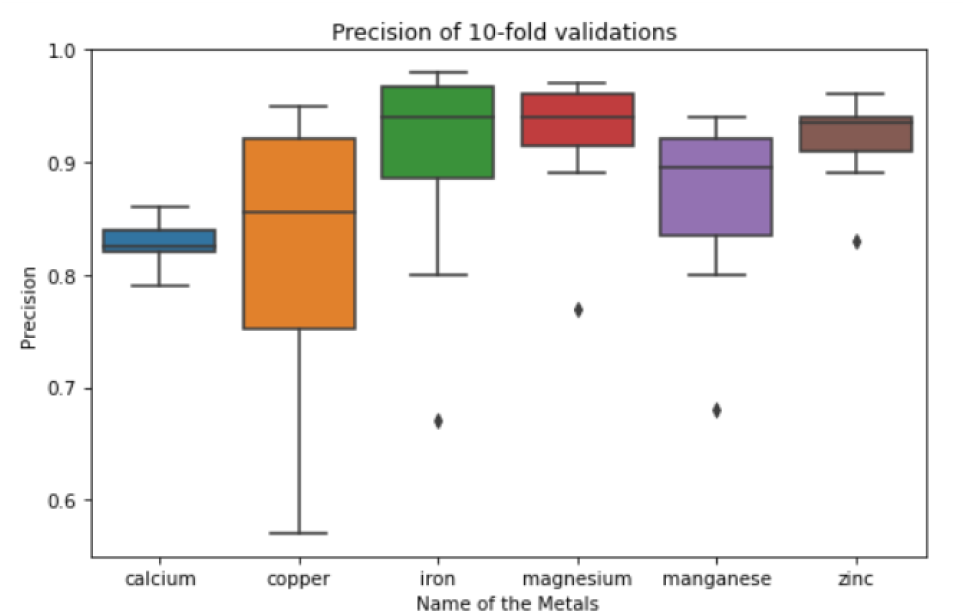
The box-plot shows the Precision of 10-fold cross validation results for 6 metals. Comparison of the metals clearly indicates that the zinc and magnesium have stable precision values of 93 *±* 3% and 91 *±* 3% respectively. The overall distribution of this two metals is not very scattered and also has low variance values. On the contrary, The copper, iron, and manganese have larger percentile values with some outliers also.

**Fig. 4.**
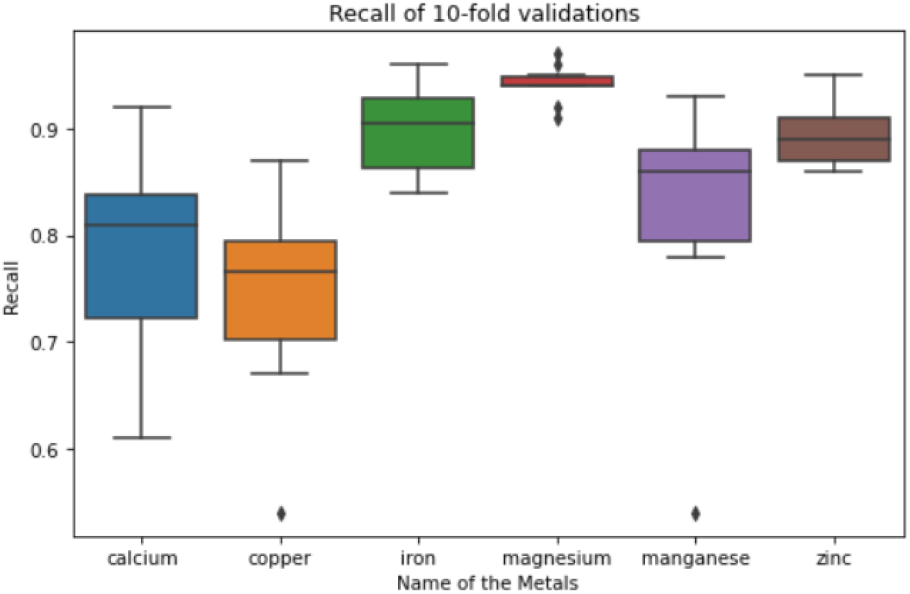
The box-plot shows the Recall of 10-fold cross validation results. Zinc has outperformed compared to other metals with a recall value of 95 *±* 2%. However, there are some outliers in that distribution due to the limitation of the samples. The calcium and copper, although having limited number of samples, performed well with a recall values of 82 *±* 5% and 78 *±* 4%.

**Fig. 5.**
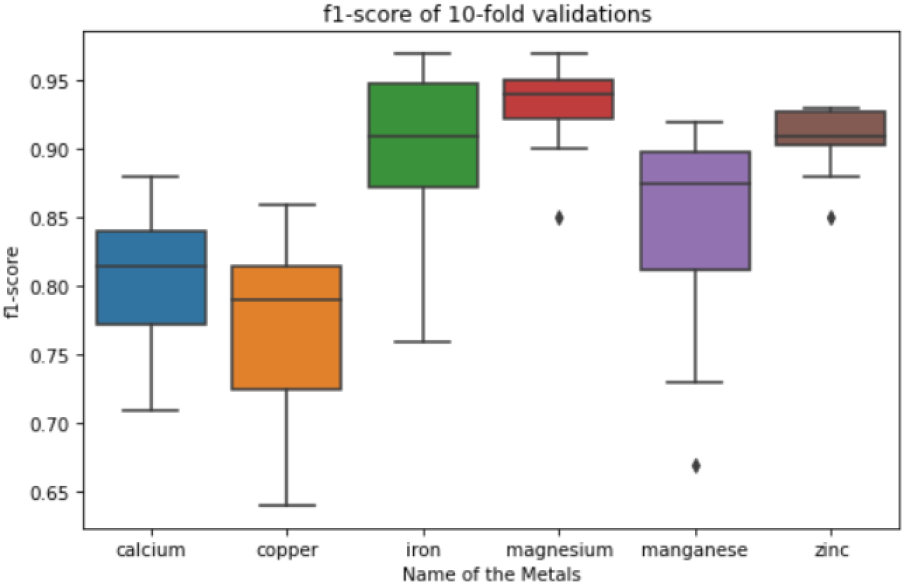
The box-plot shows the f1-score of 10-fold cross validation. The iron, magnesium, and zinc have performed well in terms of f1-score and variation is also low compare to other metals.

**Fig. 6.**
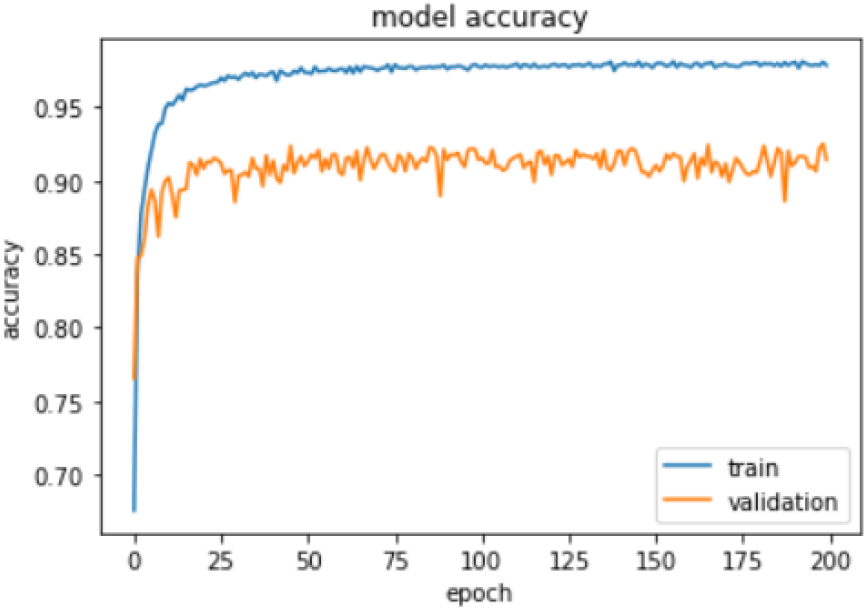
Train and validation accuracy score of the proposed model. The overall trend of validation curve remained stable over the epochs during training.

### The Impact of Positional Information in Proposed Model

It has been estimated that more than one-third of the entire proteomes are metal-binding proteins [6]. Therefore, metal position in a protein can have a significant impact on protein binding. Metal ions, such as zinc and copper, can act as co-factors in enzyme reactions and can also play a role in stabilizing protein-protein interactions. Moreover, the binding of metal ions to specific amino acid residues within a protein can affect the protein’s conformation and activity also. In some cases, metal ions can act as a “switch” in protein activity, allowing or preventing binding to other molecules. Keeping all in the mind, our experiment was designed with and without concatenating sites information with the sequences. We also tried to figure it out that whether site of a metal can contribute while making prediction or not. The network architecture was developed based-on that workflow. It was found in the experiment that the model performed better with the positional information than without providing it while making the prediction of the metals. This is also a major finding and contribution to this work.

## Conclusion

In conclusion, predicting metal binding sites in proteins is a complex and multifaceted problem that requires integrating multiple types of information and developing sophisticated computational methods. While significant progress has been made in this area, there is still much to be learned about the roles of metal ions in biological systems and the computational approaches appropriate to predict where they bind to proteins. To this end, in this paper, we have described a transformer-based model called MetaLLM to incorporate the benefits of highly sophisticated self-supervised deep learning technique to propose an end-to-end computational pipeline that is capable of predicting the presence of metal ions as well as the binding sites with state-of-art performance in terms of precision, recall and f1-score. The performance of MetaLLM is validated on a dataset of protein sequences extracted from UniPortKB/SwissProt. Being a transformer-based model, MetaLLM leverage the attention mechanism for capturing long-distance residual dependency which is crucial for identifying and distinguishing the binding sites. MetaLLM is limited to sequences in the range of 50 to 1000 amino acids. In the future, we envision to extend the model for longer sequences. Furthermore, we aim to include structural pipeline as a separate stream to validate the prediction from structural context.

## Author contributions statement

B.S. conceived the idea, B.S, F.S, F.R, S.S designed the experiment(s) and evaluation(s) plan, F.S and S.S conducted the experiment(s), B.S, F.R, S.S and F.R analysed the results, F.S, B.S, F.R and S.S wrote and reviewed the manuscript.

## Acknowledgments

The authors thank the anonymous reviewers for their valuable suggestions.

## Footnotes

1 https://github.com/DeepChainBio/bio-transformers

